# Permutation tests for hypothesis testing with animal social network data: problems and potential solutions

**DOI:** 10.1101/2020.08.02.232710

**Authors:** Damien R. Farine, Gerald G. Carter

## Abstract

1. Permutation tests are widely used to test null hypotheses with animal social network data, but suffer from high rates of type I and II error when the permutations do not properly simulate the intended null hypothesis.
2. Two common types of permutations each have limitations. Pre-network (or datastream) permutations can be used to control “nuisance effects” like spatial, temporal, or sampling biases, but only when the null hypothesis assumes random social structure. Node (or node-label) permutation tests can test null hypotheses that include nonrandom social structure, but only when nuisance effects do not shape the observed network.
3. We demonstrate one possible solution addressing these limitations: using pre-network permutations to adjust the values for each node or edge before conducting a node permutation test. We conduct a range of simulations to estimate error rates caused by confounding effects of social or non-social structure in the raw data.
4. Regressions on simulated datasets suggest that this “double permutation” approach is less likely to produce elevated error rates relative to using only node permutations, pre-network permutations, or node permutations with simple covariates, which all exhibit elevated type I errors under at least one set of simulated conditions. For example, in scenarios where type I error rates from pre-network permutation tests exceed 30%, the error rates from double permutation remain at 5%.
5. The double permutation procedure provides one potential solution to issues arising from elevated type I and type II error rates when testing null hypotheses with social network data. We also discuss alternative approaches that can provide robust inference, including fitting mixed effects models, restricted node permutations, testing multiple null hypotheses, and splitting large datasets to generate replicated networks. Finally, we highlight ways that uncertainty can be explicitly considered and carried through the analysis.

## INTRODUCTION

Permutation tests are arguably among the most useful statistical tools for the modern biologist. They are commonly used in ecology (Gotelli & Graves 1996), biogeography (Harvey 1987), community ecology (Miller, Farine & Trisos 2017), and in studies of ecological networks (Dormann *et al*. 2009) and social networks (Croft *et al*. 2011). Permutation tests randomize (or re-assign) observed data to generate a distribution of statistic values expected under a given null hypothesis. Researchers create case-specific null models by permuting data in specific ways (e.g. constraining permutations within specific groups) while keeping other aspects of the dataset the same (e.g. where and when observations were made). They are particularly useful when the standard assumptions of other statistical tests are violated, as is often the case with social network data (see Farine 2017 for a general introduction). However, several recent studies (Evans, Fisher & Silk 2020; Puga-Gonzalez, Sueur & Sosa 2021; Weiss *et al*. 2021) have highlighted issues with using social network data to test null hypotheses about the relationship between a predictor and a response (i.e. conducting regressions using network data).

The first issue is caused by the fact that different kinds of permutations create different null hypotheses. *Pre-network permutations* (or datastream permutations) are used to test how observed social network structure differs from what is expected if animals made random social decisions. This approach permutes the observed data to create many expected networks that could have occurred in the absence of any social preferences. The null hypothesis here is that, after removing the specified effects, the social structure itself is random (e.g. individuals have no other social preferences). This is a different null hypothesis than what researchers typically want when performing a correlation or regression with network data (Weiss et al. 2021). To test the statistical significance of a correlation or regression, a common permutation approach is to use *node permutations* (or node-label permutations), as used in Mantel tests and the quadratic assignment procedure (QAP) tests. By only permuting the node labels, this approach removes the statistical relationship, while preserving the same observed network properties in all expected networks. For a typical node permutation test, the null hypothesis is that there is no statistical relationship between a predictor (e.g. kinship) and response (e.g. association rate) in the *observed* network, which is correct for regression-based questions.

Drawing inferences from the observed network is, however, a challenge for animal social data. In the social sciences, the observed social network often accurately reflects affiliations, relationships, or rates of contact. In many ecological studies of animal behaviour, by contrast, the *observed* social network is typically a non-random sample that does not directly or even accurately represent the “real” social network of dyadic social preferences or contact. Instead, the observed network is typically shaped simultaneously by multiple confounding “nuisance effects”—that is, biological and methodological factors besides the hypothesized effect of interest. For example, the structure of an observed social network could be shaped primarily by individual site preferences, habitat constraints on movement, or methodological biases. An individual animal (node) may be less connected to others in the observed network only because it is harder to observe or identify, it uses a smaller subset of sampled locations, it has its main home range outside the study area, or it left the study population early. Another example is that individuals at the edge of a study area (compared to individuals at the centre) might have many associations with individuals that were never observable. For further discussion, see *Illustrating the Drivers of Type I and II Error* in Appendix.

Sampling biases will vary in magnitude and importance across study designs, but they are often inevitable. Even automated methods such as proximity sensors (Ryder *et al*. 2012; Ripperger *et al*. 2020) or barcodes (Crall *et al*. 2015; Alarcón-Nieto *et al*. 2018) are not free of sampling biases if animal-borne proximity sensors vary in their sensitivity (e.g. due to tiny differences in soldering) or if some barcodes are more difficult to identify by computer vision. Of course, unreliable observations, sampling biases, and other nuisance effects are not unique to animal social data, but these problems are especially troubling for social network analyses because observations are not independent (e.g. a single under-sampled node affects all the edges with all other nodes). The interacting roles of biological and methodological nuisance effects on the observed network structure can be difficult to identify and disentangle, and failing to do so can easily lead to spurious inferences (Farine & Aplin 2019).

Nuisance effects can be accounted for using a range of approaches. They could be estimated using covariates or random effects in a parametric model within a frequentist or Bayesian frameworks, or it is possible to control for nuisance effects using null models, for example by constraining permutations within blocks of time or space. There are many potential advantages to the parametric approach including an ability to explicitly measure the magnitude of each effect and rely less on p-values (Franks *et al*. 2021; Hart *et al*. 2021). However, as most existing statistical models of social network data have been developed in the social sciences, where observed social networks are relatively unbiased representations of the real network, it remains unclear how well—or easily— these approaches can cope with the multiple kinds of nuisance effects common in observational studies of animal behaviour. Capturing all nuisance effects in the model becomes increasingly challenging as the number of interacting nuisance effects increases, and when they do not have regular patterns across time or space. For example, one cannot simply fit individuals’ locations as a random effect by assigning individuals to a singular spatial location when home ranges are continuously distributed and overlapping in space. Further, once the network has been created, the nuisance effects generally cannot be recovered from the network itself.

The second common approach involves using pre-network permutations to simultaneously control for multiple potential nuisance effects by constraining swaps of the data to occur within blocks of time or space (Whitehead, Bejder & Ottensmeyer 2005; Whitehead 2008; Sundaresan, Fischhoff & Dushoff 2009; Spiegel *et al*. 2016; Farine 2017). In doing so, the null model can hold constant many features of the observed data including group sizes, number of observations per individual, individual variation in space use (and therefore spatial overlaps with all others), temporal auto-correlation in behaviour, temporal overlap among all pairs of individuals, the distribution of demographic classes across space and time, the variation in the density of individuals across space and time, and differences in sampling effort across space and time. Problems arise, however, when testing the relationships between a predictor and response because pre-network permutations simulate a completely different null hypothesis (i.e. random social structure outside the specified effects). As a result, the permuted networks do not have socially realistic network-level properties of real animal societies, such as the natural variance in edge weights or distribution of degree values. Therefore, when a social network has other sources of social structure beyond the effect of interest, pre-network permutation tests will produce highly elevated false positives when testing the significance of a correlation or regression (Puga-Gonzalez, Sueur & Sosa 2021; Weiss *et al*. 2021).

Our goal in this paper is to address this limitation of pre-network permutation tests, allowing them to be applied to regression while robustly accounting for multiple common nuisance effects. We propose an initial solution, which we show can maintain rates of both false positives and false negatives to around 5% under a range of scenarios. Our approach (Figure 1) uses pre-network permutations to account for nuisance effects by constraining swaps of observations within blocks of time and space, and then node permutations to conduct a nonparametric test for the effects of X on Y. This “double permutation”approach tests the null hypothesis that there is no relationship between a predictor and deviations from random social structure (within the specified temporal and spatial constraints). Specifically, we use pre-network permutations to estimate the deviation of each unit’s observed measure from its expected random value (e.g. the difference between a node’s observed degree and the median values of the same node’s degree across the permuted networks). We call this observed – expected difference the deviation score (e.g. Δdegree). These deviation scores, which account for nuisance effects, are then fit into a model of interest to generate a test statistic, and a node permutation test provides the p-value for the test statistic.

**Figure 1.**
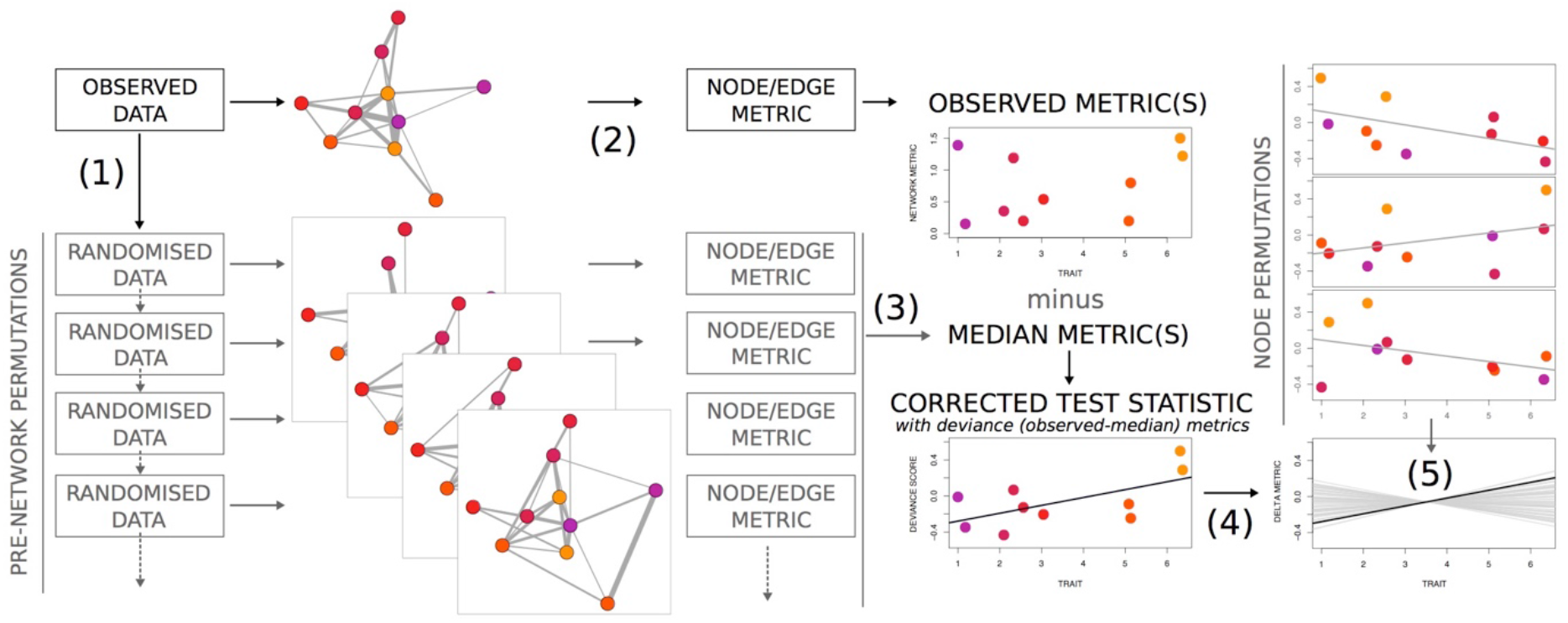
Overview of the double permutation method. First, pre-network permutations (1) generate a null distribution of expected metric values for a unit of interest (e.g. a node’s degree or an edge’s weight) alongside each unit’s observed metric (2). Next, to remove nuisance effects for each unit, the median of expected metric values is subtracted from the observed metric value, yielding a corrected metric value called the “deviation score” (3). Then, to generate a corrected test statistic, we fit a model with the deviation scores (4). To generate a p-value, a nonparametric node permutation test can then be used to compare the corrected and expected test statistics (5). Alternatively, an appropriate parametric model could be used to replace (4) and (5).

Using deviation scores is conceptually similar to generalised affiliation indices (Whitehead & James 2015), but, rather than fitting covariates in a regression model, the deviance of each unit’s metric from random expectations are extracted from pre-network permutations. Similar to extracting residuals from a model, this approach subtracts a measure of central tendency of the randomised data from the observed data, because no single permuted network can be considered as the true expected random network. As a measure of central tendency for the expected values, we use the median because the distributions of expected values are often highly non-normal and long-tailed. However, use of the median assumes that the expected values have a unimodal distribution. To check this assumption and other properties such as symmetry, we recommend visualizing the distribution of expected values generated by the pre-network permutation (see *Potential Improvements* in the Appendix).

We call this a “double permutation” because we enter the deviation scores extracted from the pre-network permutations into a node-label permutation test (e.g. QAP), but one could also opt to enter the deviation scores into an appropriate parametric model. Indeed, the basic procedure can be applied to any model for calculating test statistics, such as those generated from regressions, Mantel tests (Mantel 1967), network regression models like MRQAP (Dekker, Krackhardt & Snijders 2007), and metrics such as the assortativity coefficient (Farine 2014). We also show that the double permutation approach performs well with ‘gambit-of-the-group’ association data and with data collected using focal observations. We acknowledge that this is only one potential and imperfect solution, and we therefore also highlight alternative hypothesis-testing methods that are worth evaluating further.

## TESTING THE ROBUSTNESS OF THE DOUBLE PERMUTATION APPROACH

We evaluate the performance of the double permutation tests when using gambit-of-the-group data (simulation 1), focal sampling data (simulation 2), and dyadic observations (simulation 3), both in the absence of any real relationship and when there are strong nuisance effects. In simulations 1 and 2, we simulate the process of testing for a relationship between an individual trait and a social network metric, using three common weighted metrics (see Farine & Whitehead 2015): weighted degree (or strength), eigenvector centrality, and betweenness. In simulation 3, we simulate the process of testing for a link between pairwise kinship and association rate in a species that exhibits other social preferences not based on kinship. For each simulated dataset, we calculate p-values using several tests: node permutations, node permutations in which the model controls for covariates (number of observations and, where possible, location), pre-network permutations, pre-network permutations on the t-statistic (the equivalent of scaling the predictions, as proposed by Weiss *et al*. 2021), pre-network permutations in which the model controls for covariates (number of observations and, where possible, location), and the double permutation method. We create the networks and conduct the permutation tests using the R package *asnipe* (Farine 2013) and use the package *sna* (Butts 2008) to calculate network metrics.

### Simulation 1: Regression between a node metric and a trait using gambit of the group data

We first simulate a researcher using group-based observations to test whether individuals with a given trait value are more gregarious under three biological scenarios: (i) individuals choose groups at random, (ii) individual choice is predicted by a social trait (*T_s_*), whereby individuals with higher trait values choose to be in larger groups, and (iii) individual choice is predicted by a spatial preference (*L_s_*), whereby individuals with higher trait values are not more social but prefer sites with more resources that also hold more individuals. The last scenario represents an alternative driver of gregariousness that is challenging to control in model-fitting or using node permutations by location because individual spatial preferences cannot be reduced down to a single value. Such variation in spatial distribution of individuals is common, for example the number of great tits (*Parus major*) in Wytham Woods varied ten-fold across different parts of the woodland (Farine *et al*. 2015). In each scenario, we simulated 100 replications for varying combinations of network size (5 to 120 individuals) and mean numbers of observations per individual (5 to 40). For each simulation (one network), we extract β coefficients (slopes) and t statistics by fitting the model, weighted metric ^~^ trait (+ covariates where applicable), using the lm function in R. Note that pre-network permutations inherently control for the number of observations of each individual, and we also control for where observations were made by restricting swaps to within locations. Further details for simulation 1 are in the Appendix.

### Simulation 2: Estimating differences in group means using focal sampling data

We next simulate a researcher using focal sampling data to test whether different classes of individuals vary in their gregariousness. We build on the code of Puga-Gonzalez, Sueur and Sosa (2021) and Farine (2017), whereby each simulation assigns individuals to observations, with each observation having a focal individual and connections made between the focal and each individual observed. Simulations can be run with and without a sex difference in gregariousness. When the difference is present, females are made more gregarious by being disproportionately present in observations containing many individuals. The simulations can also introduce an observation bias, whereby females are often not observed even when present, whereas males are always observed when present. Such biases are common in field studies—for example in a study on vulturine guineafowl (*Acryllium vulturinum*) (Papageorgiou *et al*. 2019), juveniles are marked on the right wing and are only identifiable if their right side is observed, whereas adults are marked with leg bands that can be identified from any direction. The code runs 500 simulations for each scenario (sex effect or not, observation bias present or not), with parameter values that are randomly drawn from uniform distributions as follows: population size ranging from 10 to 100, observation bias ranging from 0.5 to 1 (where 1 is always observed), the female sex ratio ranging from 0.2 to 0.8, and the number of focal follows ranging from 100 to 2000. The simulation procedure was designed to allow an estimate of false positive rates (when no effect should be present but one is detected) and false negative rates (when an effect is present, but masked by the observation bias, and therefore not detected), and of the effect sizes before and after the observation bias occurs. For each simulation (one network), we extract β coefficients and t statistics by fitting the model, weighted metric ^~^ sex (+ covariates), in the lm function in R. Simulation 2 has been used in several studies to estimate performance of permutation approaches (Farine 2017; Puga-Gonzalez, Sueur & Sosa 2021) as it can capture both the pre- and post-biased metrics.

### Simulation 3: Regression between edge weight and a dyadic relationship such as kinship using dyadic observations

Finally, we simulate a researcher testing for a link between kinship and social network structure, where edges are association rates that are driven by both kinship and other social preferences. To evaluate the impact of other social preferences on error rates, we generate simulated networks in which individuals have three types of social associates: (i) weak associates, (ii) preferred associates, and (iii) strongly-bonded associates. Our simulations comprise two scenarios. In the first, association rates (edge weights) are independent of kinship, which we achieve by randomizing the kinship matrix after generating it. In the second, associations are kin-biased such that strongly-bonded associates have higher kinship on average and weak associates and non-associates (missing edges) have on average lower kinship. For each type of social network and scenario, we created ‘real’ social networks and then create ‘observed’ social networks by simulating observations. We simulate 100 replications for combinations of network sizes, with 5 to 120 individuals and the mean numbers of observations per individual ranging from 5 to 40. For each simulation (one network), we conduct a multiple regression quadratic assignment procedure (MRQAP) with the model: edge weight ^~^ kinship (+ covariates). Further details for simulation 3 are in the Appendix.

## THE DOUBLE PERMUTATION APPROACH IS ROBUST TO TYPE I AND TYPE II ERRORS

Our simulations confirm that the double permutation method is robust to type I errors across a range of scenarios. In simulation 1 (Table 1), pre-network permutation tests suffer from elevated false positives (type I error rate of 26%), consistent with previous studies (Evans, Fisher & Silk 2020; Puga-Gonzalez, Sueur & Sosa 2021; Weiss *et al*. 2021). We further show that the tendency for pre-network permutation tests to generate type I errors is greater in smaller networks and when more data are collected (Figure S2, see also Evans, Fisher & Silk 2020). Pre-network permutations using the t value as a test statistic are also prone to false positives, and perform relatively poorly at detecting a true effect (detecting them consistently less often than other approaches). Combining pre-network permutation tests together with model structures that also control for nuisance effects, when estimating the test statistic, also appears to produce unreliable results. As expected, node permutation tests perform particularly poorly when the effects are driven by non-social nuisance effects, such as variation in the spatial distribution of individuals, even when controlling for individuals’ locations in the model. The parametric p-values produced similar results to the node permutations. When node permutations were applied to a more informed model, the p-values were not clearly better or worse.

**Table 1.**
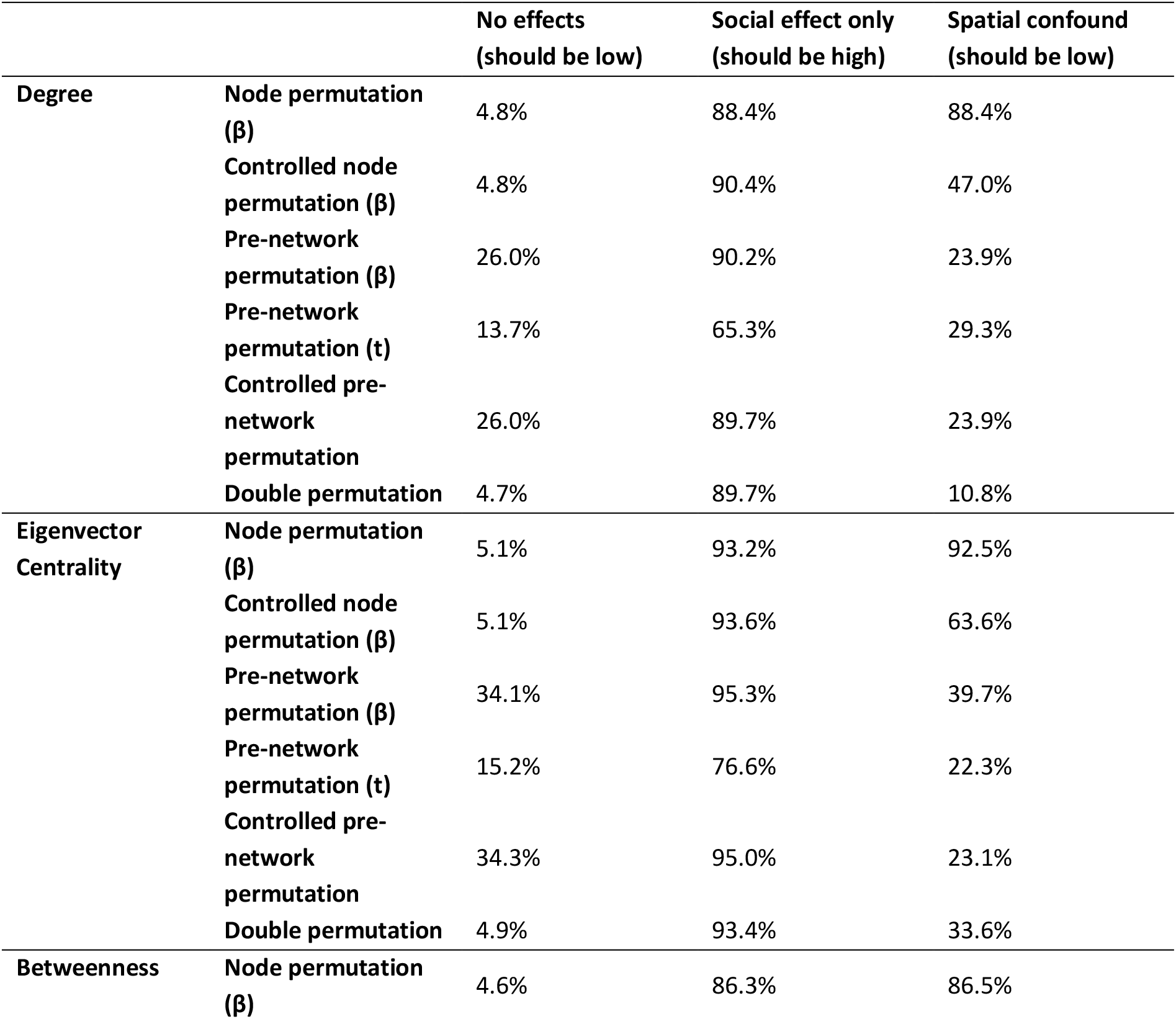
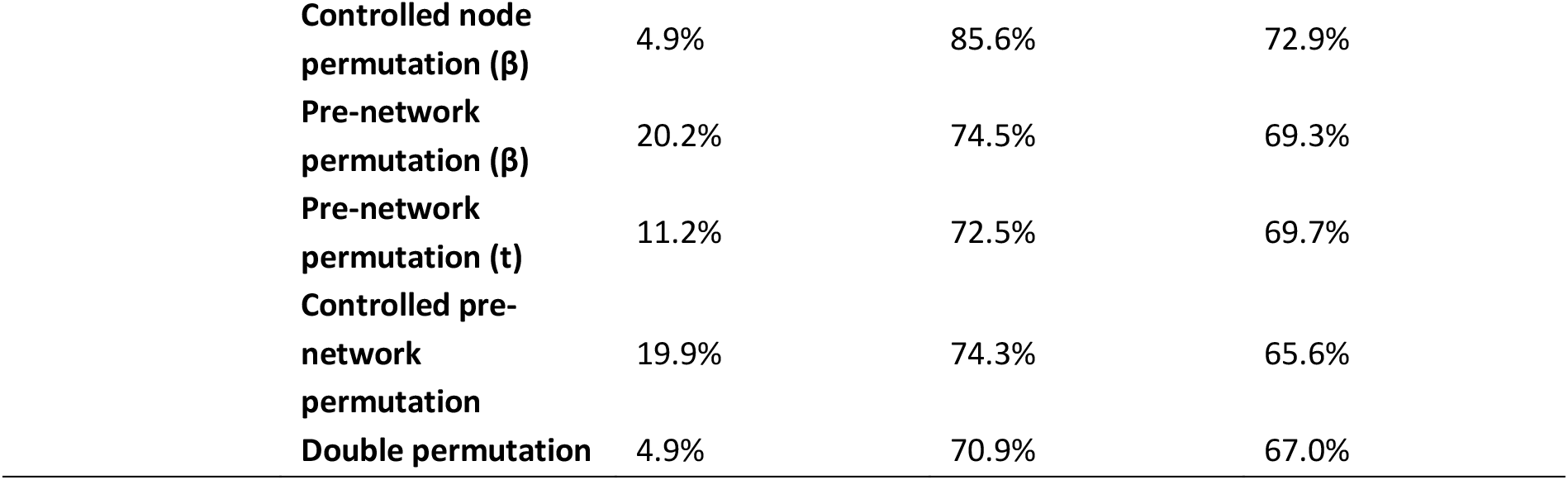
Propensity for different permutation tests to yield errors or detect real effects when using regression models to test hypotheses on networks collected using gambit-of-the-group data (model 1). Table shows the proportion of statistically significant results for an effect of a trait on degree under three sets of scenarios. When no effects are present, the expected proportion of significant results should be 5%. When a social effect is present, most results should be significant. When a spatial confound is present, the proportion of significant results should again approach 5%. Controlled node permutations include the number of observations in the models plus individuals’ most common location in the spatial confound condition. Supplementary Figures S2–S4 show how the proportion of significant results is affected by the number of observations and the number of nodes in the network.

|  |  | No effects<br>(should be low) | Social effect only<br>(should be high) | Spatial confound<br>(should be low) |
| --- | --- | --- | --- | --- |
| Degree | Node permutation ($\beta$) | 4.8% | 88.4% | 88.4% |
| | Controlled node permutation ($\beta$) | 4.8% | 90.4% | 47.0% |
| | Pre-network permutation ($\beta$) | 26.0% | 90.2% | 23.9% |
|  | Pre-network permutation (t) | 13.7% | 65.3% | 29.3% |
|  | Controlled pre-network permutation | 26.0% | 89.7% | 23.9% |
|  | Double permutation | 4.7% | 89.7% | 10.8% |
| Eigenvector Centrality | Node permutation ($\beta$) | 5.1% | 93.2% | 92.5% |
| | Controlled node permutation ($\beta$) | 5.1% | 93.6% | 63.6% |
| | Pre-network permutation ($\beta$) | 34.1% | 95.3% | 39.7% |
|  | Pre-network permutation (t) | 15.2% | 76.6% | 22.3% |
|  | Controlled pre-network permutation | 34.3% | 95.0% | 23.1% |
|  | Double permutation | 4.9% | 93.4% | 33.6% |
| Betweenness | Node permutation ($\beta$) | 4.6% | 86.3% | 86.5% |

**Table 1. Propensity for different permutation tests to yield errors or detect real effects when using regression models to test hypotheses on networks collected using gambit-of-the-group data (model 1).**
|  |  |  |  |
| --- | --- | --- | --- |
| **Controlled node permutation ($\beta$)** | 4.9% | 85.6% | 72.9% |
| **Pre-network permutation ($\beta$)** | 20.2% | 74.5% | 69.3% |
| **Pre-network permutation (t)** | 11.2% | 72.5% | 69.7% |
| **Controlled pre-network permutation** | 19.9% | 74.3% | 65.6% |
| **Double permutation** | 4.9% | 70.9% | 67.0% |

The double permutation test performs generally better than the other methods we tested across the three scenarios, producing conservative p-values when no effect is present, detecting the relationship between a trait and social metrics when present, and being more conservative than other tests when the effect is driven by non-social factors.

All tests appeared to struggle with confounded measures of eigenvector centrality and betweenness, but this is likely driven by the simulations not always producing datasets in which eigenvector centrality was strongly linked to spatial location, thereby largely over-estimating the false positive rate. However, because all methods were tested on identical simulated datasets, their relative performance can still be meaningfully compared.

In simulation 2 (Table 2), pre-network permutations again show elevated type I error rates in the absence of a true difference between classes of individuals and node permutations again show elevated type I error rates in the presence of nuisance effects. Controlling for nuisance effects in the model with node permutations helps under some circumstances (when there is a strong effect and an observation bias), but not others (e.g. when there is no effect and a bias, or when combined with pre-network permutation tests). The double permutation test almost always performs as expected. One exception is the high levels of type II errors for betweenness. This may occur because betweenness is an unstable metric (i.e. adding one edge can substantially change the distribution of betweenness values in the whole network) and pre-network permutations are therefore not generating a meaningful null distribution on which the node permutation can operate.

**Table 2.** Propensity for permutation tests to produce type I and type II errors from datasets simulating focal sampling (model 2). Simulations comprise four scenarios: (1) females and males have identical social phenotypes and are observed equally, (2) females are more social and both sexes are observed equally, (3) females and males have identical social phenotypes but observations are biased towards males (20% of observations of females are missed), and (4) females are more social but observations are biased towards males (20% of observations of females are missed).

|  |  | No observation bias |  | Observation bias ('nuisance' effect) |  |
| --- | --- | --- | --- | --- | --- |
|  |  | Phenotypes equal (Type I errors) | Females more social (Type II errors) | Phenotypes equal (Type I errors) | Females more social (Type II errors) |
| **Degree** | **Node permutation ($\beta$)** | 5.0% | 1.2% | 60.4% | 37.8% |
|  | **Controlled node permutation ($\beta$)** | 5.6% | 20.0% | 68.2% | 10.4% |
|  | **Pre-network permutation ($\beta$)** | 37.8% | 5.2% | 34.2% | 7.4% |
|  | **Pre-network permutation (t)** | 30.8% | 69.0% | 60.0% | 36.2% |
|  | **Controlled pre-network permutation** | 39.0% | 24.2% | 69.6% | 12.4% |
|  | **Double permutation** | 5.2% | 6.6% | 7.0% | 6.0% |
| **Eigenvector centrality** | **Node permutation ($\beta$)** | 4.8% | 2.4% | 55.0% | 31.6% |
|  | **Controlled node permutation ($\beta$)** | 7.0% | 2.4% | 71.4% | 7.0% |
|  | **Pre-network permutation ($\beta$)** | 45.2% | 1.2% | 43.0% | 3.0% |
|  | **Pre-network permutation (t)** | 23.4% | 74.4% | 64.4% | 34.6% |
|  | **Controlled pre-network permutation** | 38.8% | 14.0% | 79.2% | 5.2% |
|  | **Double permutation** | 4.8% | 4.2% | 6.2% | 7.0% |
| **Betweenness** | **Node permutation ($\beta$)** | 5.8% | 8.6% | 67.0% | 52.8% |
|  | **Controlled node permutation ($\beta$)** | 4.6% | 34.4% | 33.6% | 93.2% |
|  | **Pre-network permutation ($\beta$)** | 17.0% | 69.2% | 19.0% | 84.6% |
|  | **Pre-network permutation (t)** | 15.9% | 68.7% | 18.3% | 79.9% |
|  | **Controlled pre-network permutation** | 17.2% | 69.4% | 19.8% | 82.0% |
|  | **Double permutation** | 4.6% | 86.2% | 5.2% | 26.2% |

Simulation 3 (Table 3) shows that the double permutation test can reliably test null hypotheses that assume nonrandom social structure (similar to a node permutation test like QAP), such as whether association rates are predicted by kinship in cases where kinship effects are present or absent. As with the above two simulations, the double permutation test performs well when no real effect is present (i.e. type I error rates were close to 5%). All models in this simulation have elevated type II error rates because not all simulated networks with the added effect actually resulted in data with a clear effect, but the double permutation test performs more conservatively than node permutations (producing more type II errors).

**Table 3.** Propensity for permutation tests to produce type I and type II errors regarding kinship effects from simulated datasets with confounding social effects, i.e. nonrandom social structure (model 3). Table shows the type I error rates in simulations where the social effect is a confound (i.e. strong associations are not linked to kinship), and estimated type II error rates in simulations where the social effect corresponds to the hypothesis being tested (i.e. strong associations are linked to kinship). Figures S5–S6 show how the proportion of significant results is affected by the number of observations and the number of nodes in the network. While pre-network permutations appear to outperform other approaches with respect to Type II errors, this is likely because they are also more sensitive to weak effects in small networks, which are likely to correspond to type I errors rather than correctly identifying a true effect (see Figure S6).

|  | **Kinship ≠**<br>**Associations**<br>**(Type I errors)** | **Kinship ∝**<br>**Associations**<br>**(Type II errors)** |
| --- | --- | --- |
| **Node permutation ($\beta$)** | 5.1% | 12.5% |
| **Controlled node permutation ($\beta$)** | 4.9% | 12.0% |
| Pre-network permutation ($\beta$) | 18.6% | 6.2% |
| Pre-network permutation ($t$) | 3.0% | 79.1% |
| Double permutation | 5.2% | 16.5% |

## RECOMMENDATIONS

Our results confirm that pre-network permutations by themselves cannot be used to generate a p-value for a correlation, regression, or any comparison of means measured from the observed network. Having an understanding of the system and data collection procedure, and considering the possible nuisance effects that might create differences between the observed and actual network, is always critical. If the observed network is indeed an accurate reflection of the real network, then node permutations, the double permutations test described here, or any well-specified parametric models can be used (how to do this is beyond the scope of this paper).

Our results suggest that across scenarios, the double permutation test often performs better at testing null hypotheses using regressions than the other approaches when there are strong confounds, such as sampling biases or spatial preference drivers. These create high rates of false positives in tests that do not include pre-network permutations. However, the double permutation test can be too conservative when applied to unstable global node metrics such as betweenness (Table 2), which might be better studied using node permutations (in the absence of confounding effects). Because the suitability of permutation methods has not been exhaustively tested with measures of betweenness, we cannot recommend any solution for hypothesis testing with betweenness in the presence of nuisance effects. Our recommendations for choice of permutation test across node metrics and scenarios are summarised in Table 4.

**Table 4.** Recommendations from simulations for choice of permutation tests. In the absence of nuisance effects, or when weak nuisance effects can effectively be controlled in a node permutation (e.g. using restricted node permutations), then either node permutations or double permutations are likely to provide robust inference for most local network metrics (e.g. degree, eigenvector centrality) or for relational data (tests on edge weights). In the presence of nuisance effects, double permutations tests are generally recommended, except for betweenness.

|  | Metric |  |  |  |
| --- | --- | --- | --- | --- |
|  | Degree<br>Centrality | Eigenvector<br>Centrality | Betweenness<br>Centrality | Relational<br>(edge weight) |
| No clear nuisance effects | node or double | node or double | node | node or double |
| Nuisance effects expected | double | double | *unclear* | double |

Despite our focus on permutation tests, one should not infer from this study that permutation tests (or other nonparametric tests) are necessarily superior to parametric models. Carefully fitting a parametric model (or a generative model) that can explicitly capture all effects (both effects of interest and nuisance effects), identified through careful inspections of residuals and other model diagnostics, can have many benefits (see alternative approaches). For better or worse, one benefit of nonparametric tests is that they avoid the process of having to specify the best possible statistical model to fit the data. A permutation test simply compares observed and expected model coefficients (in this case extracted from the lm and MRQAP functions in R) as descriptive metrics rather than as interpretable parameters of biological meaning. However, this limits inference to null hypothesis-testing, rather than estimation of the magnitude and confidence of effect sizes (Franks *et al*. 2021).

## THE CHALLENGE OF CALCULATING EFFECT SIZES WITH SAMPLING BIAS

Inference will always benefit from relying less on p-values and instead focusing more on estimating effect sizes (Nakagawa & Cuthill 2007), and the same is true for network data (Webber, Schneider & Vander Wal 2020). Permutation tests do not, in and of themselves, provide interpretable effect sizes, meaning that the interpretation of the biology arising from the test is limited to the p-value. Franks *et al*. (2021) show that, under certain conditions, the coefficients of regression models applied to network data can generate reliable relative effect sizes after controlling for the number of observations. Multiple confounded nuisance effects can create more difficult problems when using models to estimate effect sizes. We explore this using simulation 2, where one category of individuals is both more social and less observable. Such a situation is common in nature, for example female birds that are more drab than males (e.g. most sexually dimorphic birds), shyer birds that are more social and more difficult to detect (Aplin *et al*. 2013), subadults that are more central but more difficult to recognise than adults (e.g. because they have fewer markings or as in the guineafowl example cited above). Gregariousness and detectability can be linked if individuals vary in their habitat preferences; for example, cooperatively breeding striated thornbills *Acanthiza lineata* are more social than the brightly coloured scarlet robins *Petroica boodang* that they flock with, but also more arboreal and camouflaged resulting in fewer observations per individual (Farine & Milburn 2013).

In simulation 2, the original coefficient (before the observation bias) and estimated coefficient (with the observation bias) are correlated (r=0.54), yet controlling for the number of observations of each individual consistently inflates the estimated coefficient size (Figure S7). We then test whether regression models can recover the original coefficient value using two approaches to fitting the number of observations as a covariate. First, we use a naïve model, where the scaled number of observations is simply added as a covariate. Second, we use a more informed model where the number of observations is also added as an interaction with the effect of sex (since exploration of the data would show that the number of observations differs between the sexes). As expected, the naïve model performs worse, producing estimated effect sizes that are on average 1.8 times the original value (and up to 5.1 times the original value), but correctly fitting the interaction term does not dramatically improve the estimate, with the average estimated coefficient values being 1.7 times the original value (and up to 3.3 times the original). These two models also perform very poorly at estimating effect sizes when the true effect is not present (the estimated effect sizes were on average over 250 times the true values, Figure S8). Less reliable effect sizes will also lead to less reliable p-value and hence inferences.

Nuisance effects do not need to be accounted for both within the regression model and again within the null model. Indeed, simulations show that controlling twice for nuisance effects using both covariates and permutations yields less reliable results (see results for Controlled pre-network permutation in Tables 1 and 2). In summary, a more complex model could yield correct effect sizes, but correctly accounting for multiple nuisance effects within the model is non-trivial. We encourage future work exploring this topic.

One reason why simple regression models struggle to generate robust effect sizes when controlling for nuisance effects might be because they do not deal well with individuals that are observed in groups, rather than in pairs, which is common in animal social network studies (Sah, Mendez & Bansal 2019), and known to cause problems for analysis (Evans, Fisher & Silk 2020). For such gambit-of-the-group observations, the loss of each observation can result in a variable number of edges being removed within groups. For example, when using the simple association index to estimate proportion of time spent together (Hoppitt & Farine 2018), an individual A in a group of 10 that is imperfectly observed will have a reduction in degree of 0.9 for every 10% of decrease in its detection rate, while other perfectly detected members of the group will all decrease in degree by only 0.1 per 10% decrease in the detection rate of individual A. This difference in the effects of imperfect detections on degree among members of the same group become greater as group size increase, and is not resolved by increasing sampling. Given our findings, approaches to estimating corrected effect sizes should be carefully tested before being used. Estimating effect sizes in the presence of bias is a major priority in the continued development of robust tools for animal social network analysis.

## ALTERNATIVE APPROACHES

While the double permutation test performs similarly, and usually better, than the single permutation procedures across a range of scenarios, many alternative approaches or methodological refinements can improve the robustness of inferences from hypothesis testing. Here we discuss some alternative and further approaches, but also note that not all of these methods have been exhaustively tested for performance.

### Non-permutation approaches

While this paper is focused on addressing existing problems with nonparametric permutation tests, there are also compelling arguments for moving away from permutation tests (Franks et al. 2021, Hart et al. 2021). Hart *et al*. (2021) argue that, for network regression analyses, well-specified parametric models should be used in favour of permutation tests because mixed (or hierarchical) models and Bayesian approaches can account for nonindependence in the data (e.g. by including actors and receivers as random effects), regressions tested with node permutations can violate the exchangeability of residuals, and because parametric or Bayesian approaches will reduce or remove emphasis on p-values and instead increase focus on estimating effect sizes. It would therefore be helpful to have more studies testing if and how parametric models can reliably control for common types of sampling biases and other nuisance effects described in this paper. Ideally, such studies should compare the performance of different model-fitting and permutation procedures across a range of ecologically realistic data collection scenarios, and also assess the severity of the problem of nuisance effects under various forms of data collection.

One study, Gimenez *et al*. (2019) proposed to deal with sampling biases by using capture-recapture models to explicitly model heterogeneity in detections, thereby providing more accurate estimates of network metrics. Studies estimating phenotypic variance using animal models have also proposed methods to decompose multiple sources driving between-individual variation in trait values (Thomson *et al*. 2018). Such multi-matrix models have recently been applied to animal social networks as a means of identifying the relative importance of different predictors in driving differences in social network metrics (Albery *et al*. 2021).

### Restricted node permutations

For many studies, it may be sufficient to use node permutations and control for nuisance effects by restricting which individuals’ data are swapped when performing the randomization. Such restricted node permutations are useful if individuals can be easily allocated to a distinct spatial location, or if there are clear categories of individuals that correspond to biases. Say, for example, that individual animals enter the study in distinct waves because of a standard dispersal time or because a study expanded at some point to include new individuals. In this case, permutations could be restricted to only allow swaps among individuals that entered the study at approximately the same time. However, if multiple nuisance effects have to be accounted for, one can rapidly run out of sets of exchangeable individuals. For example, a study with 40 individuals that aims to restrict swaps by two parameters (e.g. age and location) would have on average only 10 individuals per class if both parameters are binary, only 6-7 per class if one is trinary, and only 4-5 per class if both are trinary.

### Study-specific simulations

How can we be sure that a chosen method is effective? One approach is to explicitly test how sensitive a given dataset is to generating false positives or false negatives under different hypothesis-testing approaches. The procedure we used here—simulating a random trait value for each node in a network and running through the full hypothesis-testing procedure—can be a straightforward way of characterising the robustness of any given study’s results. This procedure simply involves generating a random trait variable (e.g. drawing a trait value from a normal distribution) and testing how this value corresponds to the metric of interest from the observed network using the same code as for the real variable(s) being studied. By repeating this procedure many times, one can observe the proportion of the tests that incorrectly produce a significant p-value. This result can be also reported as evidence of the selected method’s reliability. It is worth exploring further how this study-specific information might be used. For example, one might be able to correct the threshold for rejecting the null hypothesis to the point where the expected false positive rate will be 5%. Shuffling the actual node values and running a pre-network permutation test (and repeating these two steps many times) might provide an even more precise estimation of the true false positive rate, as it will be fully conditioned on the real observation data.

### Multiple null models

Testing one null hypothesis can encourage confirmation bias, and might only reject a strawman hypothesis of little interest (e.g. that chimpanzees have random interactions). Strong inference (Platt 1964) therefore requires considering and testing multiple alternative hypotheses. Similarly, multiple null models can be used to collectively examine the different processes that might be shaping the patterns present in observation data, and can be highly informative. Multiple null models are particularly effective for generating an understanding of the effects of social decisions versus space use on social network structure (e.g. Figure S1). While it is important to control for the contribution of ‘nuisance’ spatial effects to social network structure when testing hypotheses about social decision-making, the process by which animals use space (and its links to social structure) is itself an important biological question (Webber & Vander Wal 2018; He, Maldonado-Chaparro & Farine 2019). We show an example of this in Figure S1, where both social and spatial processes shape the differences in the social connectedness of males and females. Pre-network permutations that control for space would discard the biological drivers of space use (and, consequently, group size) as a nuisance effect. Aplin *et al*. (2015) evaluated the extent that the spatial distribution of individuals contributed towards their repeatability in social network metrics by reporting the distribution of repeatability values from a spatially-constrained permutation test. Farine *et al*. (2015) used two different permutation tests to identify the expected effects of individuals choosing social groups versus choosing habitats. Implementing multiple null models requires careful consideration of elevation in false positives (Webber, Schneider & Vander Wal 2020). Looking forward, the practice of developing multiple null (or reference) models using permutation can be further extended to include generative (e.g. agent-based) models that test specific alternative hypotheses (Hobson *et al*. in press).

### Bootstrapping (and its limitations)

Another approach that is often considered to be useful for estimating uncertainty (e.g. confidence interval around effect sizes) is bootstrapping (Lusseau, Whitehead & Gero 2008; Farine & Strandburg-Peshkin 2015; Bonnell & Vilette 2021), which involves resampling the observed data with replacement to create new datasets of the same size as the original. This procedure can estimate the range of values that a given statistic can take, and whether the estimate overlaps with an expected null value (see Puth, Neuhauser & Ruxton 2015). Bootstrapping, however, is not always appropriate as a means of hypothesis testing in animal social networks, because like node permutations, it relies on resampling the observed data under the assumption that the observed network reflects the true social structure. For example, missing edges in an association network represent an association rate of zero, but in many cases these zero values could actually be weak associations that exist in the real world but were not observed. In such cases, bootstrapping the edge weights suggests that these unobserved edges have no uncertainty, which is obviously false. Thus, bootstrapping social network data should only be used with care.

### Replicated networks

Pre-network permutation tests were initially designed to evaluate whether the social structure of a population is nonrandom, given sparse association data (Bejder, Fletcher & Brager 1998), and error rates for regression-based hypotheses degrade as more observations are collected for a few nodes (Figure S2). The good news is that many observations allow the creation of replicated networks (Hobson, Avery & Wright 2013), which is achieved by splitting the dataset to produce multiple networks (without overlapping observations). When doing so, it is important that each replicated network contains sufficient data to produce reliable estimates of network structure (Farine 2018). The same hypothesis-testing procedure can then be applied to each network independently. Using emerging methods for automated tracking, social networks can be created for each season (e.g. Papageorgiou *et al*. 2019), across periods of several days (e.g. Dakin *et al*. 2021), each day (e.g. Boogert, Farine & Spencer 2014), or even each second (e.g. Blonder & Dornhaus 2011). Independent networks that produce consistent results when tested independently provides much stronger support for a given hypothesis than any single network can. If, instead, effects are unstable over time, this might suggest either the presence of other underlying dynamics that warrant further investigation, more careful analyses, or the need for longer time periods for each replication.

Any inference becomes stronger again if each of the replicate social networks contains different sets of individuals or if the network is reformed in each time period. An example of this are within-roost association networks that are formed each day after foraging bats all leave and come back into the roost before sunrise (Ripperger *et al*. 2019). Alternatively, a hypothesis could be tested on alternative subsets of the populations, such as different communities (Bond *et al*. 2020; Bond *et al*. 2021). Given sufficient data, replicated networks could be combined by using tools from meta-analyses to estimate an overall effect size. However, such an approach would need to ensure that the same biases don’t impact each of the networks in the same way.

Although within-study replication can improve our confidence in a given result, ultimately the gold standard is replication across studies, as within-study replications cannot control for many of the nuances in how data are collected, stored, and analysed. One example of a replication study tested the effects of developmental conditions on the social network position of juvenile zebra finches (*Taeniopygia guttata*). In the original study (Boogert, Farine & Spencer 2014), birds were given either stress hormone or control treatments as nestlings, and their social relationships were studied after they became nutritionally independent from their parents. In the replication study (Brandl *et al*.2019), clutch sizes of wild zebra finches were manipulated to experimentally increase or decrease sibling competition (a source of developmental stress), and social associations (in the wild) were recorded after birds fledged. Across both studies, 9 out of 10 hypothesized effects had the same result (i.e. both statistically significant or not), and all 10 of the hypothesised effects were in the same direction (binomial P<0.0001).

## CONCLUSIONS

In this paper, we propose an approach that can avoids the elevated false positives that occur from pre-network permutations assuming random network structure under the null hypothesis and node permutations assuming an unbiased observed network under the null hypothesis. Our proposed solution, or the use of permutation tests more generally, does not negate the need to carefully consider statistical issues that have been highlighted for more orthodox statistical practices (Forstmeier, Wagenmakers & Parker 2017). For example, the common practice of using models for both data exploration and hypothesis testing are estimated to produce rates of type I error as high as 40% (Forstmeier & Schielzeth 2011). High false discovery rates can also be caused by overfitting, ignoring model assumptions, or choosing an incorrect model structure (e.g. by failing to fit random slopes to a mixed effects model, see Schielzeth & Forstmeier 2009). The same pitfalls occur when using these methods to calculate a test statistic for use with a permutation test. In general, false positive rates are likely to increase with the complexity of the question and the dataset, and dealing with empirical datasets in the biological sciences often requires making complex decisions for which the solutions aren’t clear—such as whether to log-transform count data and use a simple general linear model (Ives 2015) or to fit a more complex generalized linear model (O’Hara & Kotze 2010). In some cases, researchers may opt for a less powerful tool that may be easier to wield correctly.

In the context of null hypothesis-testing with social network data, each permutation procedure (including constricting the swaps in different ways) creates a specific null model (see Figure S1 for an example), so the crucial statistical consideration is ensuring that the correct null hypothesis is being tested. One particularly important point highlighted by this work, and that of others (e.g. Franks *et al*. 2021), is the need to pay close attention to the importance of different processes for a given hypothesis of interest. For example, if we assume that space use is constrained by non-social factors, and if we aim to understand animal social decisions, then space use (and its consequences) could be considered a nuisance effect. However, if we aim to study the transmission of information or pathogens, then space use is an important factor of interest contributing to the outcome of the transmission process. Thus, a given process might represent a nuisance effect for one question but not another, even if these factors represent two halves of the same feedback loop, such as habitat choices driving social preferences and vice versa (Cantor *et al*. 2021).

Permutation tests can control for a large range of nuisance effects within the null model without explicitly identifying, measuring, and controlling for every source of bias. There are both pros and cons of this approach, but we believe that the strength and robustness of permutation tests lies in their flexibility and simplicity. By using a range of permutation tests to measure the relative deviations of the observed data from different null expectations, it is also possible to evaluate the relative contribution of different processes shaping a network. Regardless of the method used, inference with social network data requires thinking carefully about what specific processes may have produced the patterns in a given observed dataset.

## ACKNOWLEDGEMENTS

We thank the Farine and Carter labs, Jordan Hart, Josh Firth, Matt Silk, Michael Weiss, Dan Franks, Lauren Brent, Sebastian Sosa, and the many researchers who contacted the authors with questions about the pre-print, for helpful comments, insights, and discussions regarding the issues presented in this paper. DRF was funded by the Max Planck Society, an Eccellenza Professorship Grant of the Swiss National Science Foundation (Grant Number PCEFP3_187058), and a grant from the European Research Council (ERC) under the European Union’s Horizon 2020 research and innovation programme (grant agreement No. 850859). DRF received additional support from the Deutsche Forschungsgemeinschaft (DFG, German Research Foundation) under Germany’s Excellence Strategy – EXC 2117 – 422037984. Work by GGC is supported by a grant from the National Science Foundation (IOS-2015928).

## AUTHOR CONTRIBUTIONS

DRF and GGC developed the method, and wrote the manuscript together. DRF developed and ran the simulations.

## APPENDIX

### ILLUSTRATING THE DRIVERS OF TYPE I AND TYPE II ERRORS

Here we clarify the main problem by considering a simple, verbal, but concrete example of why errors can arise when using permutation tests. Imagine a study population where animals cluster for warmth each night. Clustering associations include 2-10 individuals and the group composition can change from night to night. The dataset contains a list of observed clusters, their location, time, and the individuals in those clusters that could be correctly identified. From these data, researchers generate a network describing the propensity for each dyad to be observed in the same cluster with the aim of finding out if kin are more likely to cluster: do individuals preferentially cluster with kin?

First, the researchers consider a node permutation approach (e.g. using a Mantel or MRQAP test) to correlate pairwise kinship and association. However, kin can associate more than expected by chance without kin-biased behaviors or even kin discrimination—for example if siblings are born at the same time, are limited to similar home ranges, and then disperse at around the same time. Under such a scenario, a significant result from a node permutation test would correctly support the hypothesis that the network is kin-structured, but it would be spurious support for the hypothesis that kinship is a causal driver of social associations (false positive). Alternatively, kinship-biased association could be undetected or underestimated (false negative) if younger animals are both more likely to associate with kin but also less likely to be individually marked and recorded. In the presence of sampling bias or other nuisance effects, hypotheses about the process generating an effect of kinship on association could be challenging to accurately assess using basic node permutation tests (but see Alternative Approaches section for more discussion on how node permutations can be restricted to help alleviate some of these effects).

The researchers might therefore turn to using a pre-network permutation test. To create expected outcomes for the relationship between kinship and edge weights, they repeatedly swap individual observations within sampled locations and time periods. Doing so allows them to eliminate the nonsocial nuisance effects described above, but it also assumes that no other social factors cause social network structure, because pre-network permutations assume a null hypothesis of no social effects at all. Imagine that in this species, individuals always spend about 90% of their time with only 1-3 closely bonded associates. This constraint on social structure means that every node (individual) always has a large variance in edge weights (association rates) across all its possible associates, because most association rates are zeros while a few are larger (i.e. closer to one). This variance in edge weight is much greater than expected from random associations (few zeros but also few larger values). When the researchers create randomised networks using pre-network permutations, many zero association rates are replaced with non-zero values as individuals that never co-occurred are swapped into groups together, and the individuals in the resulting random networks almost never spend 90% of their time with a few individuals (i.e. the previously strong edges become weaker). Even if association rates are not kin-biased and the close social bonds are randomly distributed with regards to kinship, the extremely nonrandom social structure of the observed network could easily lead to a false positive with regards to the effect of kinship (Weiss *et al*. 2021). If even a few strong bonds exist among kin by chance in the observed data, these strong kin bonds might not ever appear in the randomized networks. This scenario occurs because the observed network typically has much higher variance in the measure of interest (edge weight or node metric) than the corresponding random networks from pre-network permutations (Aplin *et al*. 2015; Firth *et al*. 2018; Weiss *et al*. 2021). Thus, once again, the actual effect of kinship on association is challenging to accurately evaluate in the presence of a nuisance effect (in this case social preferences unrelated to kinship).

A different potential problem with pre-network permutation tests is that they can provide overly-confident estimates of minor deviations from random. In part, this problem occurs because constructing social networks requires large numbers of observations (Langen 1996; Farine & Strandburg-Peshkin 2015; Davis, Crofoot & Farine 2018) with many repeated observations of the same individuals, and in any hypothesis test a sufficiently large number of observations can invariably produce p-values well below 0.05 even when the effect size is not biologically important.

Pre-network permutations can also suffer from incorrect inference when networks contain only a few nodes. For instance, consider a simplistic network containing only three nodes (male, male and female), and with only three observed dyadic associations, with all three being between the two males. In such a scenario, a pre-network permutation testing whether females are less connected than expected by chance would be significant at p<0.05, as the observed data is the only one of 27 combinations of the three dyadic associations that would result in the female having a degree of zero (the p-value here would approach 0.037). While any inference drawn from three observations of three individuals is obviously unreliable, the same problem can be present, and much harder to notice, in more complex analyses.

The previous example illustrates problems caused by nuisance effects when the variables are traits of edges (kinship and association). The same principles apply when testing hypotheses relating the traits of individuals. For instance, a false impression that individuals of one sex are more gregarious (higher network centrality) could be caused by sex-based differences in observability, attraction to spatially clumped resources, variation in home-range size, differential survival in different habitats, or differences in the timing of their presence or absence in the population. These issues are common to many other types of ecological studies, but their influence can be exacerbated in social network studies. Some of these effects are sampling effects, and others will only cause a problem if there’s a mismatch between the question of interest to the researcher and the question being addressed by the analysis (e.g. ‘do males prefer larger groups?’ versus ‘do males occur in larger groups?’, see Figure S1).

Although it is useful to conceptually distinguish between effects that are social versus non-social, or biological versus methodological, these can be difficult to disentangle in practice, because biological biases can influence methodological biases and social behaviours can contribute to the apparently ‘non-social’ effects. Studies could benefit from making more explicit considerations of the social decisions that contribute to spatial and temporal biases. For example, individuals’ spatial ranges could be determined by density-dependence in their decisions to settle in a location or move elsewhere. We provide some advice on how permutation tests can help uncover such effects in the future directions section.

Another example is how social behaviour can influence observability. Consider a study testing if male birds have higher degree centrality. If females tend to be found at the periphery of groups, then an observer standing close to the centre of a group might be more likely to miss observations of females and their associates, whereas the observer can always detect the associates of the (predominately male) individuals at the centre of a group, thereby introducing a sex bias in the number of observed associates. An observer standing outside the group could have the opposite bias. This simple example highlights how biases can arise even when observations are made in groups where every individual is individually-identifiable, and why the focus on detecting relationships (i.e. estimating edges) makes network studies more prone to nuisance effects than many other types of studies.

**Figure S1.**
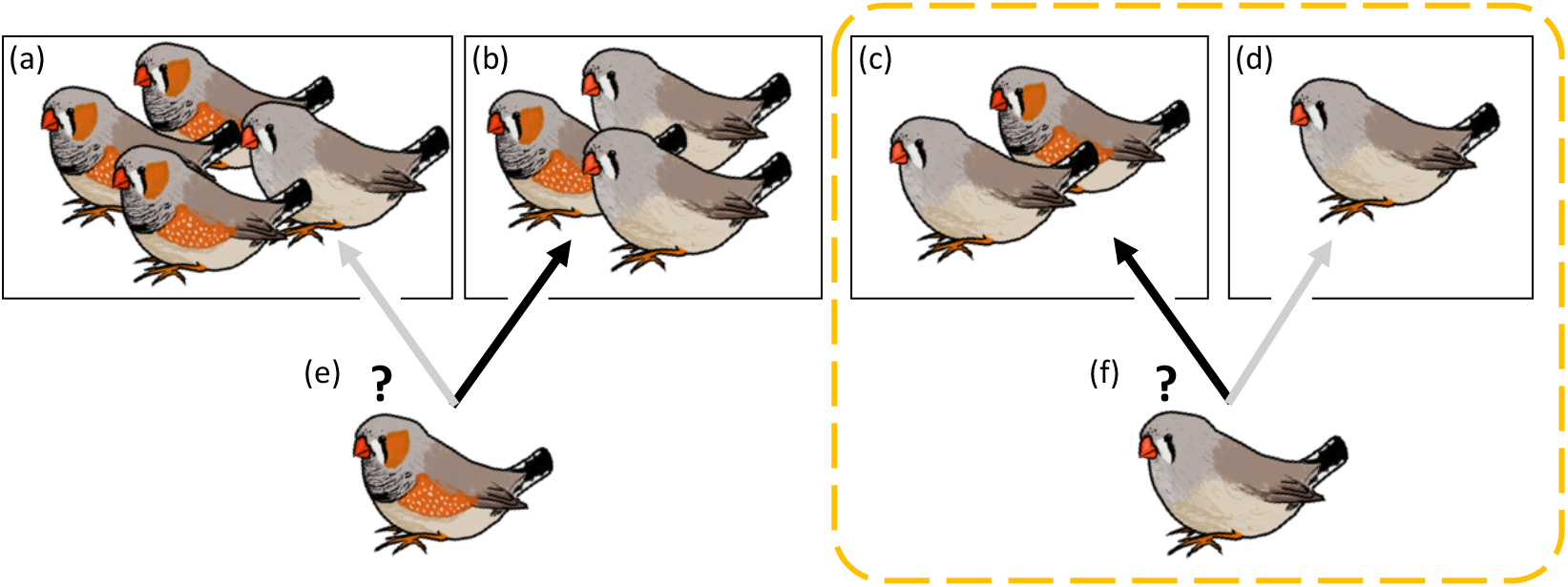
Example of the challenges associated with testing hypotheses on social data. Compared to females, are males more often in larger groups? In the scenario shown above, males (colourful individuals) are observed in larger groups (e.g. groups a, b, c) more often than are females (less colourful individuals). However, now consider a slightly different question: Compared to females, do males have a greater preference for larger groups? The same observation is not conclusive regarding the causal processes. Imagine that groups a and b are outside a territory (shown by the dashed line) defended by a single dominant male (shown in c) that allows females to enter freely and form groups (c and d) but excludes other males. Even if all males choose smaller groups (b vs a), and all females (e.g. f) choose larger groups (c vs d) given their available options, we could still observe males in larger groups than females due to the constraints imposed by the territory. Thus, even if males are equally or less gregarious, they can end up in larger groups and having more social connections. However, by permuting the individual observations within locations (a pre-network permutation), we can simulate individuals of both sexes making random social decisions given the options that were available to them. Under such a null model, a subordinate male that was never seen in the territory (groups c and d) would never be observed there in the permuted data, and would therefore always be found in larger groups. Because males would end up in larger groups in both the observed and permuted datasets, this pre-network permutation test would correctly indicate that males do not prefer larger groups. However, the same test would incorrectly fail to detect that males and females differ in their numbers of connections. The former question might be the focus of a study on how individual decisions shape network structure, whereas the latter question might be the focus of a study of how network structure shapes information or pathogen transmission.

An advantage of pre-network permutations is that they allow precise control over many possible nuisance effects, without needing to measure or even identify all of them. For example, if a lower proportion of individuals are individually-identifiable at the edge of a study area, then individuals seen at the edge would always occur in groups containing fewer individuals that can be individually-identified. In this case, controlling for space would automatically control for differences in group size that arise due to spatial variation in identification rates by only swapping individuals into other groups that also had fewer individual-identified individuals. Such an effect could be challenging to control for explicitly in a statistical model. On the other hand, pre-network permutation tests allow many other important aspects of the permuted networks to change (such as degree distributions or variances), and thus pre-network permutations alone should not be used to assess the effect of a predictor while controlling for social structure (Puga-Gonzalez, Sueur & Sosa 2021; Weiss *et al*. 2021). Ideally, one would use pre-network permutations to control for nuisance effects without making unrealistic random networks, which would include maintaining the variance in association strengths.

### POTENTIAL IMPROVEMENTS OF THE DOUBLE PERMUTATION TEST

There are many further directions and refinements that can be explored for the double-permutation test. While we have demonstrated that the method performs adequately across a range of scenarios using simulated data, it is not always possible to simulate all possible types of uncertainty that might exist in empirical datasets. For example, the median expected value might not effectively represent the value expected by the null hypothesis, because when the possible configurations of the data are severely limited (for instance by a small sample size of observations per individual), the resulting distribution of expected metric values might not be unimodal. For example, if individuals have strong group size preferences, then their expected degree might jump dramatically from their preferred values to the distribution of mean degrees from the population, as more and more swaps are made. It can therefore be useful to visualize the expected distributions of metric values across individuals when possible, and to remove individuals that have been under-sampled (Farine & Strandburg-Peshkin 2015).

Rather than using a single value (such as the median, see Figure 2), future studies could explore ways of carrying over uncertainty from the distribution of permutation values when calculating the corrected test statistic. For example, one could use a Monte Carlo approach that repeatedly samples from the distribution of permuted values when calculating the deviation score for each unit in the analysis (each node or each edge), and use these many measurements to estimate the 95% range of the corrected test statistic. Carrying uncertainty through the analysis could be implemented using a Bayesian framework (Farine & Strandburg-Peshkin 2015), or to model the dynamic process of the observation of connections in the network (Koskinen & Snijders 2007). Thus, there remains significant grounds for continued improvements in the methods for hypothesis testing using animal social network data.

For many studies, it is important to not only test a hypothesis of interest but also to accurately estimate the connection strength between individuals. One method that has been proposed are generalized affiliation indices (Whitehead & James 2015). These involve regressing the observed association strength against nuisance factors (such as home-range overlap) to generate a corrected value that accounts for the opportunity to associate. Permutation tests have also been suggested as a means of identifying nonrandom preferred or avoided relationships (Whitehead, Bejder & Ottensmeyer 2005). Yet, it remains to be determined whether permutation tests could also provide more accurate estimates of the strength of each relationship. Following these methods, it could be possible to estimate a corrected association (or interaction) strength by subtracting some measure of the distribution of permuted values from the observed value of each edge.

### SIMULATION 1 DETAILS

The first simulation starts by drawing *N* individual trait values *T_i_* from a normal distribution with a mean of 0 and a standard deviation of 2. We assign each individual to have on average K observations by drawing *K_i_* from a Poisson distribution with *λ* = *K* and balancing these values to ensure that ∑*K_i_* = *N* × *K*. We then create *G* groups, where *G* = 0.5 × *N* × *K*, and randomly assign each of these groups to have a group size value *X* ranging from 1 to 10. To allocate individuals into groups, we order the individuals from the smallest trait value to the largest to create scenarios where the trait value should impact the social behaviour of individuals (trait has social impact, *T_s_* = *TRUE*), or order these at random to create scenarios where the trait value has no relation to the social behaviour of individuals (*T_s_* = *FALSE*). We assign each individual into groups by selecting the *K_i_* groups that have empty spaces, and with a higher probability of selecting smaller groups. In doing so, individuals earlier in the order are disproportionately more likely to be assigned to smaller groups, filling them up, and leaving only larger groups for later individuals to fill, thereby creating a relationship between individuals’ trait value *T_i_* and their weighted degree *D_i_* when *T_s_* is true. Because the simulations did not always create a relationship (especially in simulations of eigenvector centrality and betweenness), we stochastically re-ordered the traits when *T_s_* = *TRUE* and no relationship was present to create a relationship. This was done by ordering individuals by their trait value and probabilistically allocating them to nodes according to the node values, thereby resulting in a strong relationship.

From these observation data, we follow the design of our double permutation method (see Figure 2) to first calculate how each node’s degree deviates from what is expected from random behaviour, and, second, to calculate the relationship between the deviance score for weighted degree values 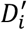 and the trait values *T_i_*. We then implement two conceptual variations, *L_s_*, to go with the two *T_s_* scenarios above. In the first variant (location effect, *L_s_* = *TRUE*), we assume that *X* corresponds to a spatial preference, such that individuals prefer patches closest to the centre of their home ranges in a one-dimensional linear environment ranging in values from 1 to 10. Patches at one end of this environment, corresponding to a larger *X*, contain more resources and therefore can hold more individuals. The second variant (no location effect, *L_s_* = *FALSE*) is a scenario in which the group size values *X* corresponds to the outcome of social decisions, such that individuals with a larger *X* have a preference for larger groups. These variants enable us to use the same code to produce a relationship between *T_i_* and network degree *D_i_* where in one scenario the decisions are social (*T_s_* = *TRUE* and *L_s_* = *FALSE*) and in another scenario the relationship between *T_i_* and *D_i_* arises from decisions that are driven by a non-social covariate (*T_s_* = *TRUE* and *L_s_* = *TRUE*). To control for the location *X_j_* that each group was observed in when *L_s_* = *TRUE*, we use within-location swaps in the pre-network permutation tests. We also calculate individual’s most common location to include as a covariate in “controlled node permutations”.

### SIMULATION 3 DETAILS

In simulation 3, we first create a ‘real’ network comprised of *N* individuals with a network density *D* drawn from a uniform distribution (ranging from 0.05 to 0.6), and selecting *Z* × *D* edges (where *Z* is the maximum possible number of undirected edges) with probabilities 0.6, 0.3, and 0.1 for edge types 1 to 3, respectively (and all other edges set to 0).

Second, we make an average of *K* observations per individual. Specifically, we create 1.2 × max(*K_i_*) sampling periods (Whitehead 2008), and randomly allocate individuals to being observed in *K_i_* of these sampling periods. To create associations for each sampling period, we then select all pairs of individuals with an edge present in the real network and where both are present in that sampling period, and draw a 0 or 1 to signify whether they were observed together or not. Here we set the binomial probability of being observed together when they are both present to be 0.1 for edges of weak associates, 0.6 for edges of preferred associates, and 0.9 for edges of strongly-bonded associates, based on the real network. We select the number of individuals and observations (*N* and *K*) using the same parameter sets as in model 1.

Third, we create a kinship network, setting the kinship level of each individual based on their edge type in the real network. Specifically, we draw relative kinship values from a beta distribution with *α* = 1 and *β* = 2 (i.e. left-skewed) for missing edges and edges of weak associates, *α* = 2 and *β* = 2 (i.e. unskewed) for edges of preferred associates, and *α* = 3 and *β* = 2 (i.e. right-skewed) for edges of strongly-bonded associates. Because beta distributions range from 0 to 1, these distributions assume that 1 corresponds to the closest relatives in the population. We use these parameters to create kinship distributions with differences in mean relative kinship (0.33, 0.5, 0.6, respectively) according to social relationship type. Note that these relative kinship values can be divided by two to create a maximum kinship value of 0.5 with no effect on the outputs of the model.

Our simulations comprise two scenarios. In the first scenario, edge weights are unrelated to kinship, which we achieve by randomizing the kinship matrix after generating it (thereby keeping the same relationship between the variance in the network edges and in the kinship matrix). In the second scenario, associations are kin-biased by keeping the kinship matrix as it was generated, thus strongly-bonded associates have the highest kinship on average and weak associates and non-associates (missing edges) have on average the lowest kinship. We create the networks, conduct the pre-network permutation tests, and conduct the regressions using the MRQAP functionality in the R package *asnipe* (Farine 2013).

## SUPPLEMENTARY FIGURES

**Figure S2.**
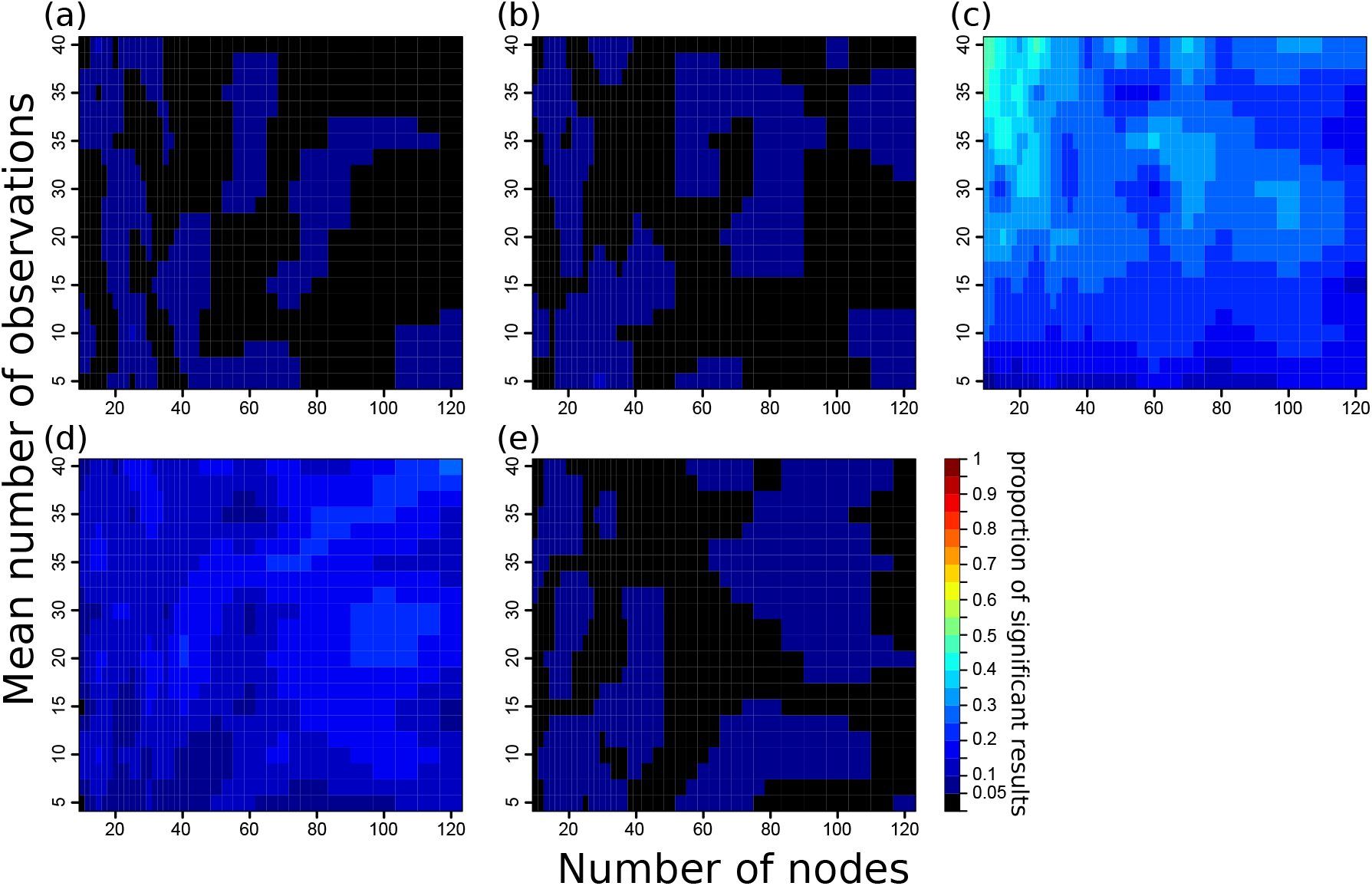
Proportions of statistically significant results for differently-sized networks and different observation efforts for simulation 1 under the scenario when *T_s_* = *FALSE* (no effect is present) and *L_s_* = *FALSE* (individuals are not preferentially located in different patches), with degree as the network metric. Panels represent the p-values calculated using (a) node permutation tests with the coefficient value as the test statistic, (b) node permutation tests controlling for number of observations, (c) pre-network permutation tests with the coefficient value as the test statistic, (d) pre-network permutation tests with the t statistic as the test statistic, and (e) double permutation tests with the coefficient as the test statistic. These results highlight the propensity for pre-network permutation tests (c) to produce spurious results when networks have few nodes but many observations (top left of the plot).

**Figure S3.**
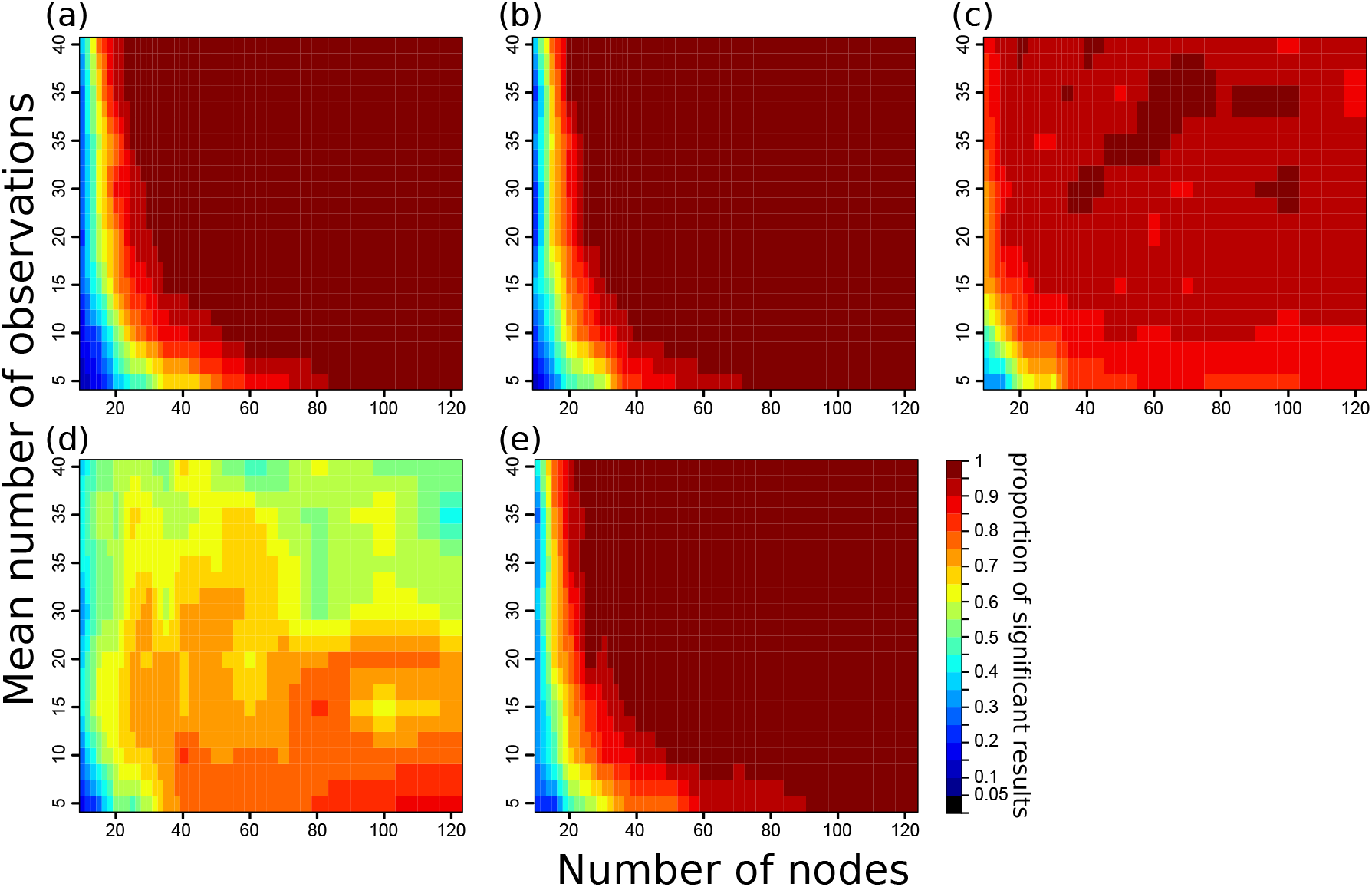
Proportions of statistically significant results for differently-sized networks and different observation efforts for simulation 1 under the scenario when *T_s_* = *TRUE* (an effect is present) and *L_s_* = *FALSE* (the effect is not a spatial confound), with degree as the network metric. Panels represent the p-values calculated using (a) node permutation tests with the coefficient value as the test statistic, (b) node permutation tests controlling for number of observations, (c) pre-network permutation tests with the coefficient value as the test statistic, (d) pre-network permutation tests with the t statistic as the test statistic, and (e) double permutation tests with the coefficient as the test statistic. These results highlight the propensity for pre-network permutation tests (c) to be more likely to produce significant results when networks have few nodes but many observations (left-hand of the plot relative to panels a, b and e). Further, results show that using the t statistic (d) produces unreliable results (i.e. the significance does not increase when more observations are made).

**Figure S4.**
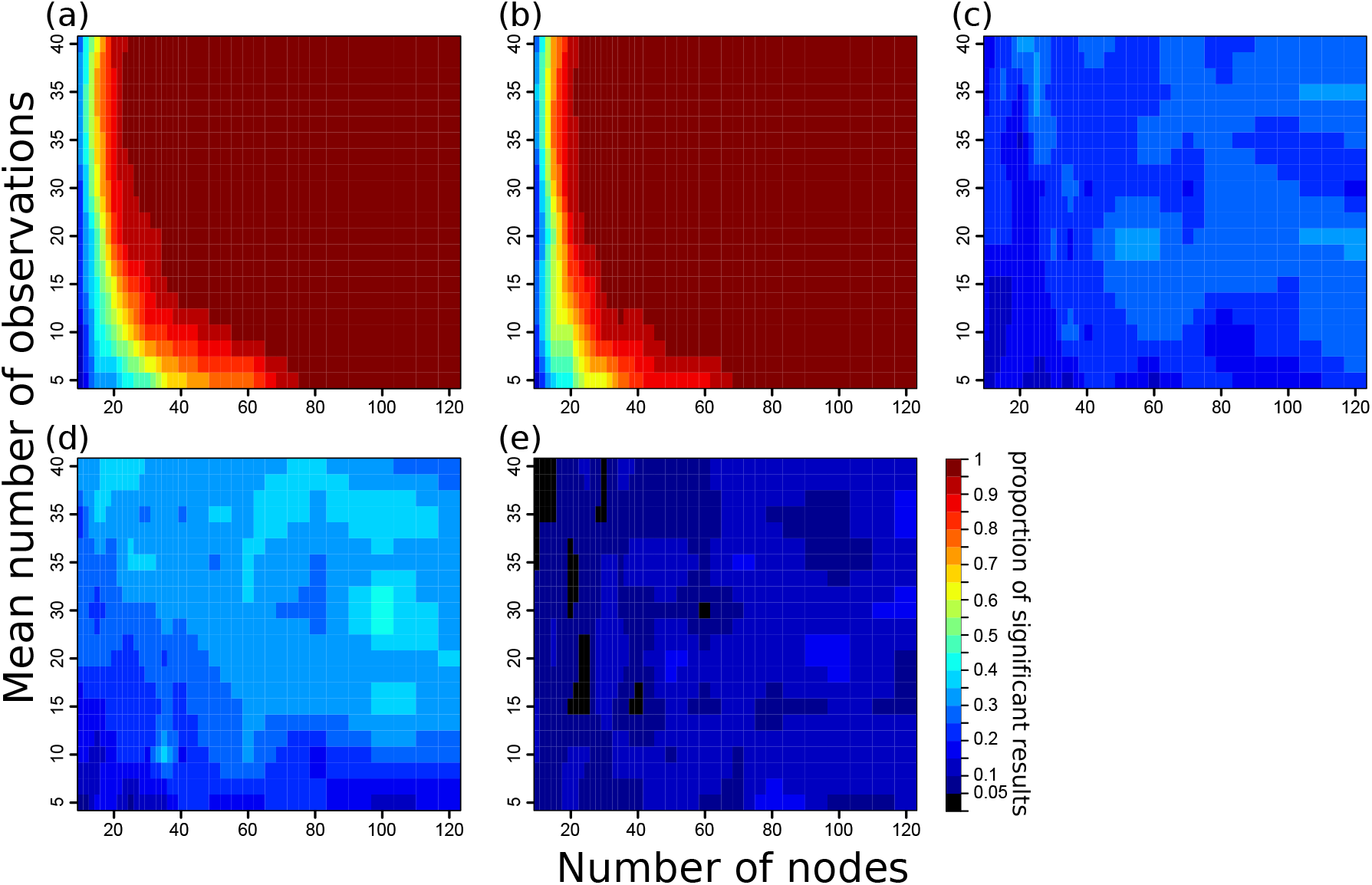
Proportions of statistically significant results for differently-sized networks and different observation efforts for simulation 1 under the scenario when *T_s_* = *TRUE* (an effect is present) and *L_s_* = *TRUE* (the effect is a spatial confound), with degree as the network metric. Panels represent the p-values calculated using (a) node permutation tests with the coefficient value as the test statistic, (b) node permutation tests controlling for number of observations, (c) pre-network permutation tests with the coefficient value as the test statistic, (d) pre-network permutation tests with the t statistic as the test statistic, and (e) double permutation tests with the coefficient as the test statistic. These results highlight the poor performance of node permutation-based models (panels a and b).

**Figure S5.**
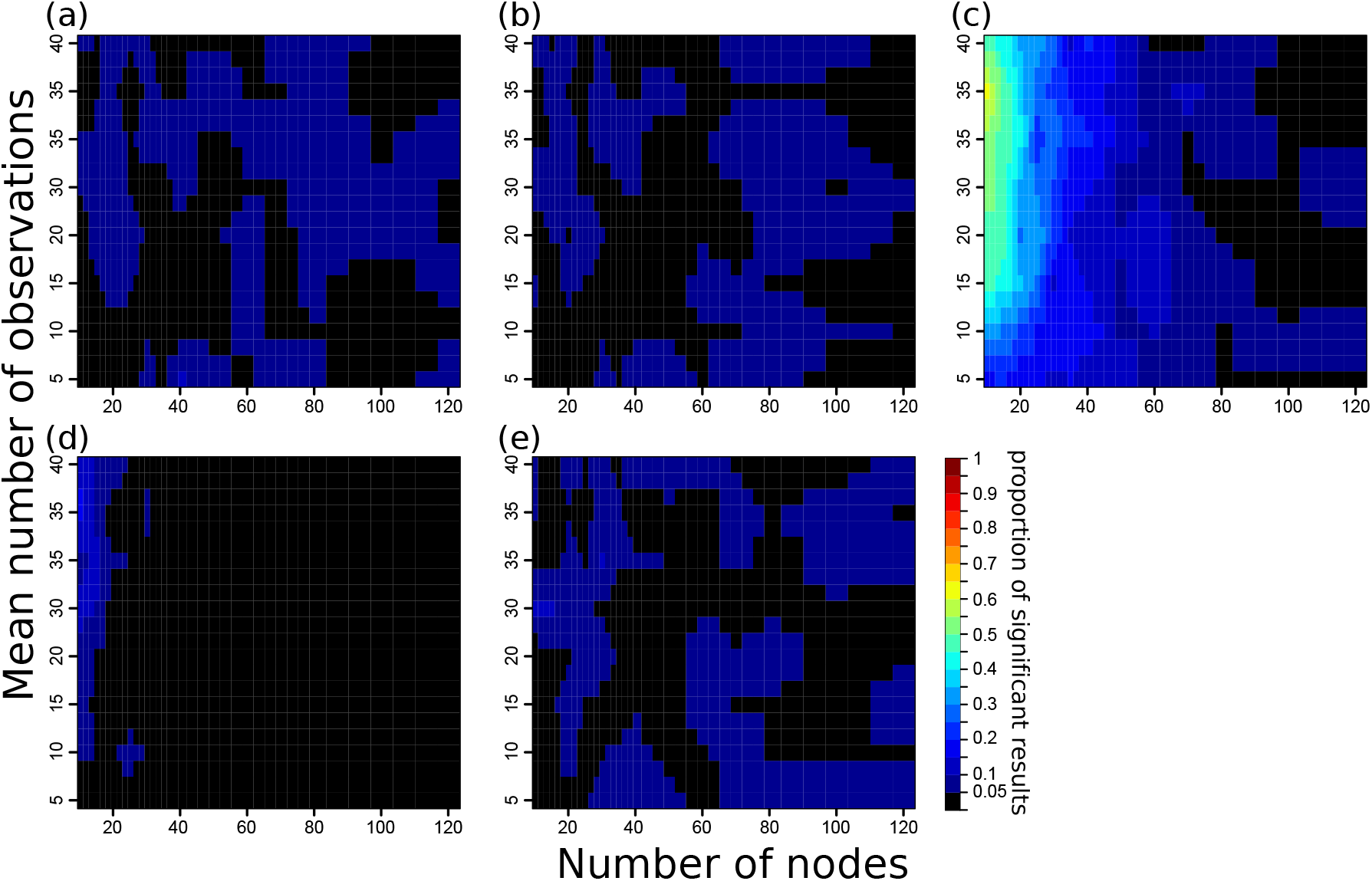
Proportions of statistically significant results for differently-sized networks and different observation efforts for simulation 3 under the scenario where kinship does not predict association rates (edge weights). Panels represent the p-values calculated using (a) node permutation tests with the coefficient value as the test statistic, (b) node permutation tests controlling for number of observations, (c) pre-network permutation tests with the coefficient value as the test statistic, (d) pre-network permutation tests with the t statistic as the test statistic, and (e) double permutation tests with the coefficient as the test statistic. These results highlight the propensity for pre-network permutation tests (c) to produce spurious results when networks have few nodes present (left-hand-side of the plot).

**Figure S6.**
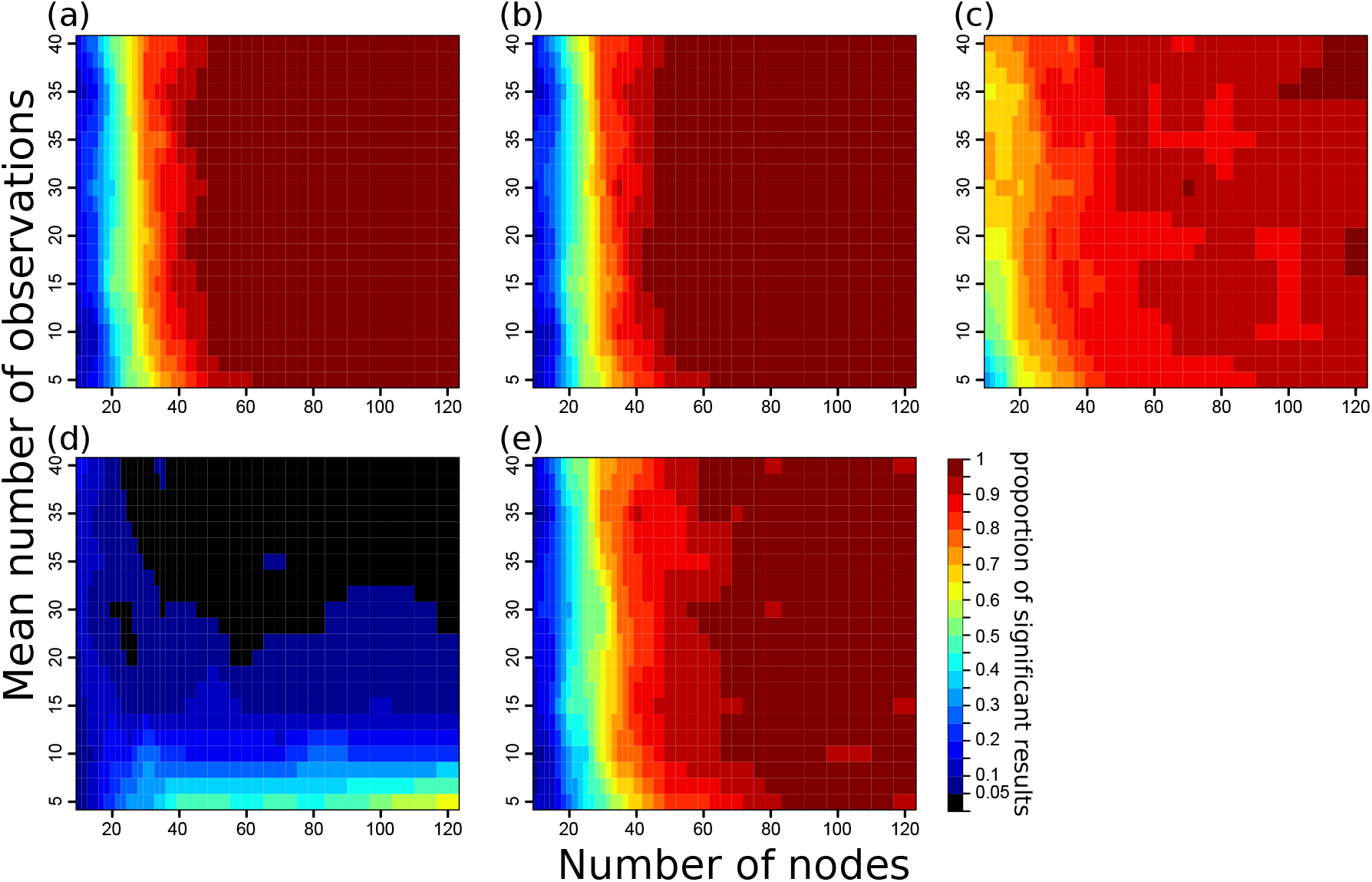
Proportions of statistically significant results for differently-sized networks and different observation efforts for simulation 3 under the scenario where kinship predicts association rates (edge weights). Panels represent the p-values calculated using (a) node permutation tests with the coefficient value as the test statistic, (b) node permutation tests controlling for number of observations, (c) pre-network permutation tests with the coefficient value as the test statistic, (d) pre-network permutation tests with the t statistic as the test statistic, and (e) double permutation tests with the coefficient as the test statistic. These results highlight the propensity for pre-network permutation tests (c) to be more likely to produce significant results when networks have few nodes but many observations (left-hand of the plot relative to panels a, b and e). Further, results show that using the t statistic (d) produces unreliable results (i.e. the significance does not increase with when more observations are made).

**Figure S7.**
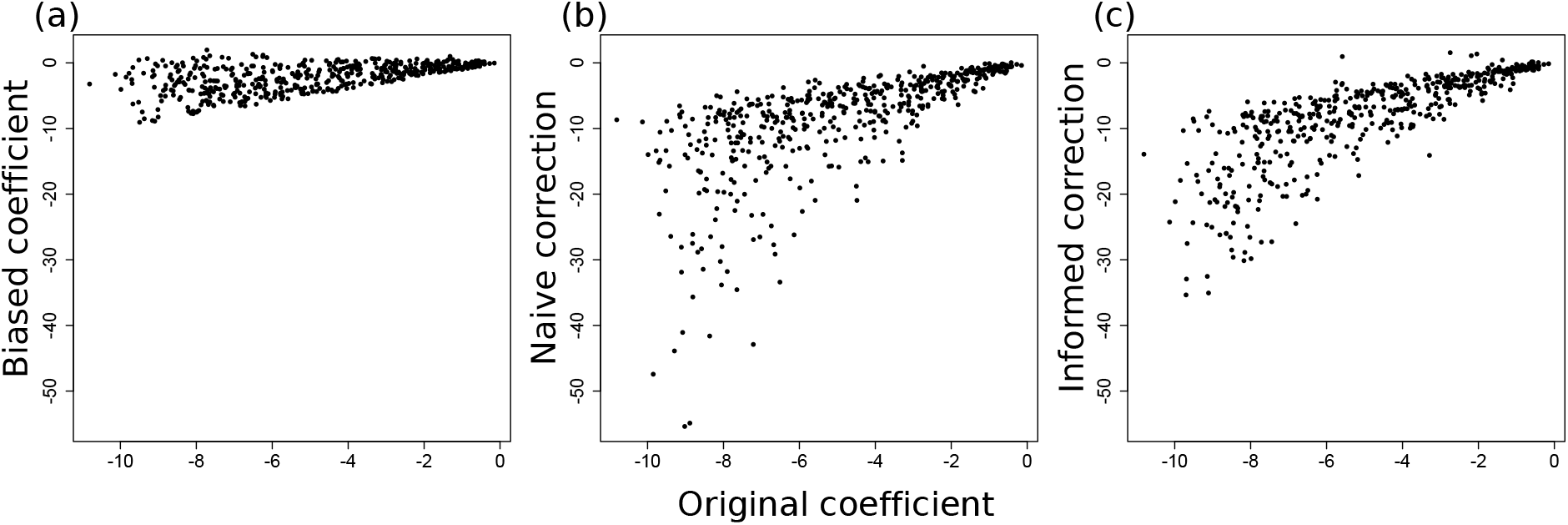
Relationship between the original coefficient value (the relationship between degree and sex prior to introducing an observation bias) and estimations of the coefficient value using the data from simulation 2 where an effect is present (females are more gregarious). (a) The original coefficient versus the coefficient estimated from the biased observations. (b) The original coefficient versus a naïve correction involving adding only the number of observation for each individual as a covariate in the model. (c) The original coefficient versus an informed correction that involves including an interaction term between sex and the number of observations. Each point represents one simulation. While these coefficients are correlated, the corrected coefficient values can be greatly over-estimated, suggesting that adding the number of observations into a model does not produce reliable effect sizes.

**Figure S8.**
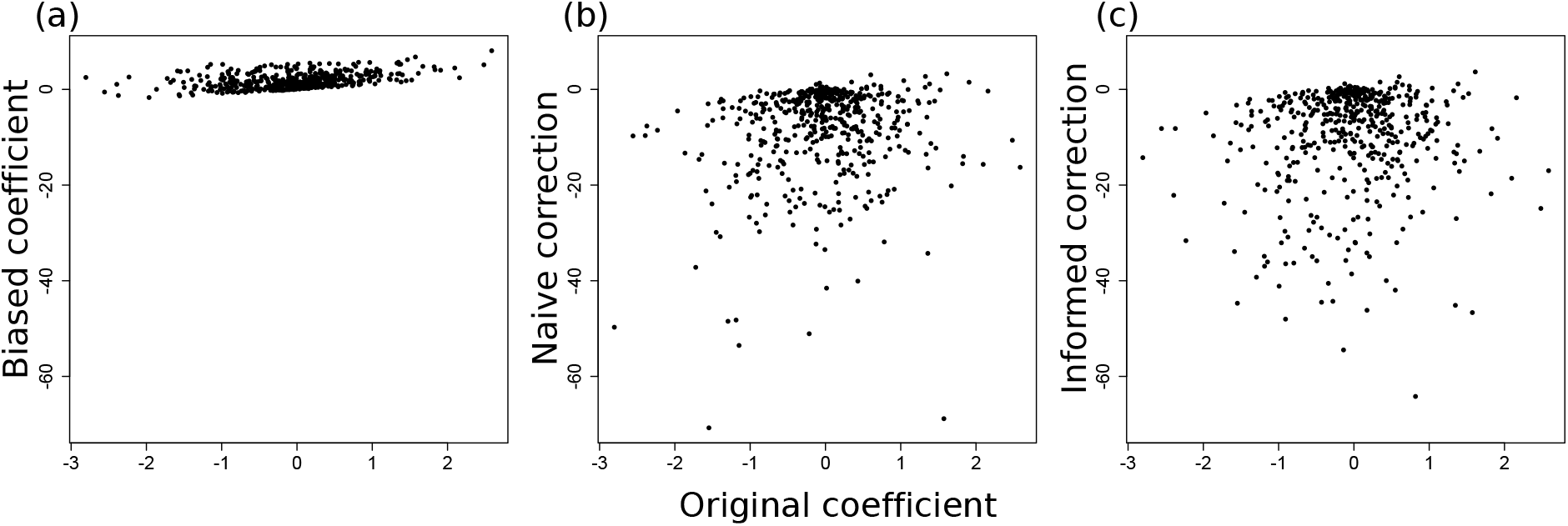
Relationship between the original coefficient value (the relationship between degree and sex prior to introducing an observation bias) and estimations of the coefficient value using the data from simulation 2 where no effect is present (females and males are equally gregarious). (a) The original coefficient versus the coefficient estimated from the biased observations. (b) The original coefficient versus a naïve correction involving adding only the number of observation for each individual as a covariate in the model. (c) The original coefficient versus an informed correction that involves including an interaction term between sex and the number of observations. Each point represents one simulation. Because there was no original effect present, the coefficients are not correlated. However, the corrected coefficient values generate extremely large coefficient values, suggesting that adding the number of observations into a model does not produce reliable effect sizes.

